# MLOsMetaDB, a meta-database to centralize the information on Liquid-liquid phase separation proteins and Membraneless organelles

**DOI:** 10.1101/2023.07.23.550222

**Authors:** Fernando Orti, María Laura Fernández, Cristina Marino-Buslje

**Affiliations:** Leloir Institute/IIBBA. Buenos Aires, Argentina; Instituto de física interdisciplinaria y aplicada (IFNIA). Universidad de Buenos Aires, Argentina; Dto. de Física. Facultad de Ciencias Exactas y Naturales. Universidad de Buenos Aires, Argentina

**Author notes:** María Laura Fernández.

## Abstract

Over the past few years, there has been a focus on proteins that create separate liquid phases in the intracellular liquid environment, known as membraneless organelles (MLO). These organelles allow for the spatiotemporal associations of macromolecules that dynamically exchange within the cellular milieu. They provide a form of compartmentalization crucial for organizing key functions in many cells. Metabolic processes and signaling pathways in both the cytoplasm and nucleus are among the functions performed by MLOs, which are facilitated by diverse combinations of proteins and nucleic acids. However, disruptions in these liquid-liquid phase separation processes (LLPS) may lead to several diseases, such as neurodegenerative disorders and cancer, among others.

To foster the study of this process and MLOs function, we present **MLOsMetaDB** (http://mlos.leloir.org.ar), a comprehensive resource of information on MLO and LLPS related proteins. Our database integrates and centralizes available information for every protein involved in MLOs, which is otherwise disseminated across a plethora of different databases.

Our manuscript outlines the development and features of MLOsMetaDB which provides an interactive and user-friendly environment with modern biological visualizations and easy and quick access to proteins based on LLPS role, MLO location, and organisms. Additionally, it offers an advanced search for making complex queries to generate customized information.

Furthermore, MLOsMetaDB provides evolutionary information by collecting the orthologs of every protein in the same database.

This is a crucial resource for transferring knowledge to other species because the LLPS protein databases are highly biased towards human proteins and, to a lesser extent, a few other well-studied or model organisms. Overall, MLOsMetaDB is a valuable resource as a starting point for researchers studying the many processes driven by LLPS proteins and membraneless organelles.

## Introduction

Membraneless organelles (**MLOs**) or biomolecular condensates are dynamic subcellular compartments that lack surrounding lipidic membranes (Banani, Lee, Hyman, & Rosen, 2017). **MLOs** are formed through a liquid–liquid phase separation phenomenon (**LLPS**) by which proteins and nucleic acid molecules self-assemble into liquid droplets maintained in a condensed phase through multivalent interactions, mostly transients and dynamics (Nott et al., 2015). The proteins or nucleic acids that play a direct role in the formation and/or the stability of the MLOs through the LLPS process, are referred to as **Drivers**. The components that do not actively participate in the formation but are present in the MLO under specific conditions, are known as **Clients**. Proteins that can alter the formation, composition, dissolution or stability of MLOs are classified as **Regulators** (Farahi, Lazar, Wodak, Tompa, & Pancsa, 2021).

Recently, several primary LLPS-dedicated databases became available with curated and not curated datasets of proteins. These databases are **PhaSePro** (Mészáros et al., 2020). **PhaSepDB** (You et al., 2020). **DrLLPS** (Ning et al., 2020) **LLPSDB** (Li et al., 2020), **RNAPhaSep** (Zhu et al., 2021) **and RSP** (Liu et al., 2021). The last two also collect RNAs involved in the LLPS process.

In a previous study by our group, it was found that current LLPS-dedicated databases differ in their focus, inclusion criterion, bias in the included protein, annotation, incompleteness, and curation levels among other characteristics, making the analysis difficult and the generation of new knowledge challenging (Orti, Navarro, Rabinovich, Wodak, & Marino-Buslje, 2021).

To address this issue, we developed **MLOsMetaDB** (http://mlos.leloir.org.ar/), a comprehensive and centralized resource of information on MLO and LLPS related proteins.

**MLOsMetaDB** integrates and centralizes protein information of the primary LLPS/MLOs databases and enriches their annotations with functional, structural and evolutionary information, focusing on molecular features associated with the LLPS process. In such a way, it concentrates available information disseminated in a plethora of different databases for every protein involved in MLOs, difficult to embrace when one or a group of proteins are the focus of a study.

Our database provides an interactive and user friendly environment, modern biological visualizations, and easy and quick access to proteins by LLPS role, MLO location and organism. It also provides an advanced search to make complex queries and generate customized information.

Also, it allows the scientific community to easily create different subsets of interest to carry out further analyzes.

Finally, it provides evolutionary information by collecting the orthologous proteins by protein in the database. This is a key resource for transferring knowledge to other species because the LLPS protein databases limitation is that they are highly enriched in human proteins and in total contain very few organisms, the most studied or model ones.

## Methods

### LLPS and MLOs Associated Proteins

Proteins were collected from four LLPS and MLOs related databases: **PhaSePro** (Mészáros et al., 2020), **PhaSepDB** (You et al., 2020), **DrLLPS** (Ning et al., 2020) **y LLPSDB** (Li et al., 2020; Ning et al., 2020). Entries were integrated and unified by Uniprot Accession, when available.

Disorder-related annotations and predictions were retrieved from MobiDB (Piovesan et al., 2021), while structural annotations were obtained from **PFAM, PDB** and **AlphaFold2 DB** (Varadi et al., 2021). Orhologs and evolutionary annotations were retrieved from **OmaDB** (Altenhoff et al., 2021).

### Biological role assessment

Proteins within the MLOs were categorized as Drivers, Clients and Regulators depending on the annotations in their source DB and/or when clear evidence was found in different datasets or literature.

#### Drivers

are proteins retrieved from PhaSePro, LLPSDB, Phase-Separation dataset from PhaSepDB and proteins annotated as Scaffolds in DrLLPS.

#### Clients

proteins retrieved from MLO dataset from PhaSepDB and proteins annotated as Clients in DrLLPS.

#### Regulators

proteins annotated as Regulators in DrLLPS.

### Data Processing

Data was downloaded and processed using Python 3.8 and Pandas package; graphs were prepared with Matplotlib 3.5.1, and Seaborn 3.5.1 libraries.

### Web Server implementation

MLOsMetaDB is stored in a MongoDB database and its backend was developed using Flask framework (Version 1.1.2). The website is implemented using VueJS and Bootstrap 5. The RestFul API and its documentation were developed using Swagger Ui Library. Interactive biological visualizations include implementations of ProSeqViewer (Bevilacqua, Paladin, Tosatto, & Piovesan, 2021), Feature Viewer (Paladin et al., 2020) and Mol* Viewer (Sehnal et al., 2021).

## Results

### Protein entries

The MLOsMetaDB dataset is composed of **12038** unique proteins obtained from the four LLPS/MLOs databases. We retrieved **121** proteins from PhaSePro, **373** from LLPSDB, **5191** from PhaSepDB and **9251** from DrLLPS (**Figure 1A**). From the total, **8365** proteins are annotated in one database and **2504** and **113** proteins in two and three databases respectively. It is worth noting that the overlap between DBs is very small, Only **56** proteins of the dataset are present in the four databases, highlighting the importance of this effort to centralize the data (**Figure 1B**).

**Figure 1.**
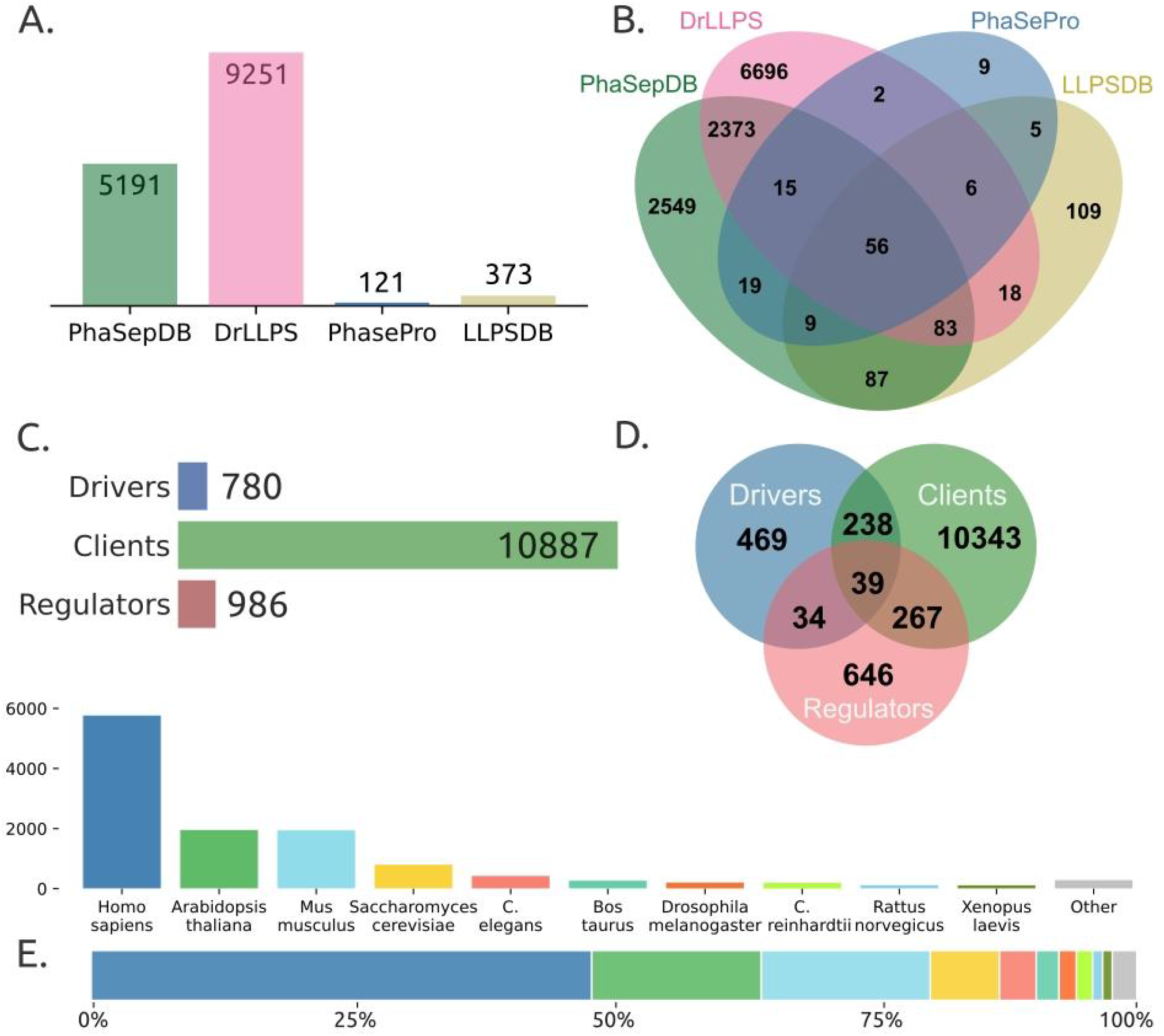
MLOsMetaDB Statistics. A) Number of proteins in MLOsMetaDB categorized by their source database. B) Venn diagram illustrating protein overlap between databases. C) Number of proteins in MLOsMetaDB classified by LLPS role annotation. D) Venn diagram showing protein overlap between LLPS roles. E) Distribution of proteins by organism: number (top) and percentage (bottom).

The imbalance in the number of annotated proteins between the different databases can be explained by the different inclusion and curation criteria of each database. PhaSePro and LLPSDB only collect manually curated proteins with experimental evidence of being LLPS Drivers while DrLLPS and PhaSepDB also include proteins from High-Throughput experiments with no evidence of LLPS behavior.

MLOsMetaDB contains annotations of **780** LLPS Driver proteins, **10887** Clients and **986** Regulators. There are **238** proteins annotated as Driver and Client, **34** as Driver and Regulator and **267** as Client and Regulator, 39 proteins are annotated as Driver, Client and Regulators (see **Figure 1C and 1D**). The overlap between LLPS roles may be due to the fact that a protein can have different roles in different MLOs or be miss classified in different databases. For instance, the human protein PCNT (Uniprot Acc: O95613) is annotated as a client protein within the centrosome in DrLLPS, whereas PhaSepDB designates it as a driver protein within the same MLO. Another example are proteins annotated with different roles in different organelles. **Figure 1 D** illustrates the organisms distribution in the DB,

Considering the biology of MLOs, it is to be expected that there will be a greater number of client proteins recruited by MLOs than of driver proteins. To caracterize a protein as driver a stronger experimental evidence is required, therefore, some proteins that are currently annotated as clients may in the future be reclassified as drivers.

### Organisms distribution

Direct experimental data is available for a limited number of proteins and there is a strong bias towards the most studied proteins and organisms. Human proteins are 47%, while the rest are distributed among few organisms, all of them well studied or model ones (**Figure 1E**). This is a limiting point since data for less popular organisms or proteins is still lacking. Having experimental support is labor intensive, time consuming and costly, so for some proteins it is not possible to have this information unless transferring it by homology.

Instead, by gathering orthologous proteins, starting from **12038** we reach **118433** other proteins in **2408** different organisms, in such a way, enriching them with valuable information to better understand their biological roles.

### Web Usage

MLOsMetaDB has been designed to facilitate users research and navigation.

The basic search is with the Uniprot accession or Gene Name. Also, in the **main page** there are quick accesses to different datasets such as a particular MLO, LLPS role and the principal model organisms. The **advanced search** allows more complex queries filtering by protein identifiers, annotation type, PFAM domains, LLPS roles, LLPS source databases and molecular features related to the LLPS process and all the possible combination of filters (As an example: mouse driver proteins, present in SG and Cajal Bodies with 30% of disorder content), among other possibilities (**Figure 2**).

**Figure 2.**
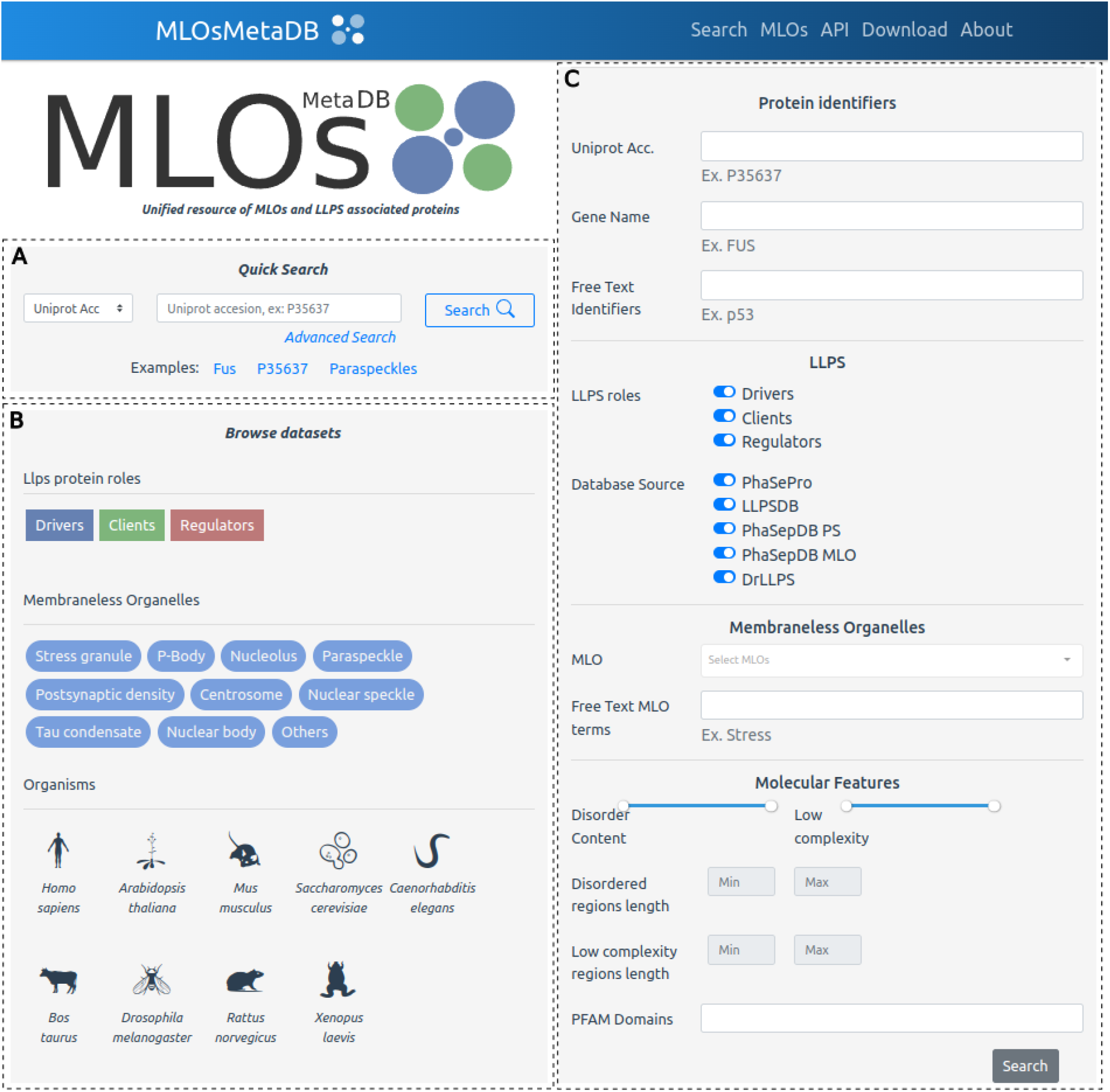
MLOsMetaDB’s Main page. A) Quick search by Uniprot Acc, Gene Name, and MLO. B) Browse datasets by LLPS role, MLO and model organisms. C) Advanced search: In this section users can realize more complex searches, combining different fields, including LLPS roles, LLPS source databases, PFAM domains, one or more membraneless organelles (exact or free text search). Also the sequence percent of disorder and low complexity content and lengths of the segments can be specified.

The **results page** offers a quick and nice view of the general features of a protein (LLPS Rol, Organism, Molecular features, IDRs, LCRs, Databases Sources) (**Figure 3)**. Results can be displayed in cards or table mode, filtered by columns and can be downloaded as JSON or TSV formats.

**Figure 3.**
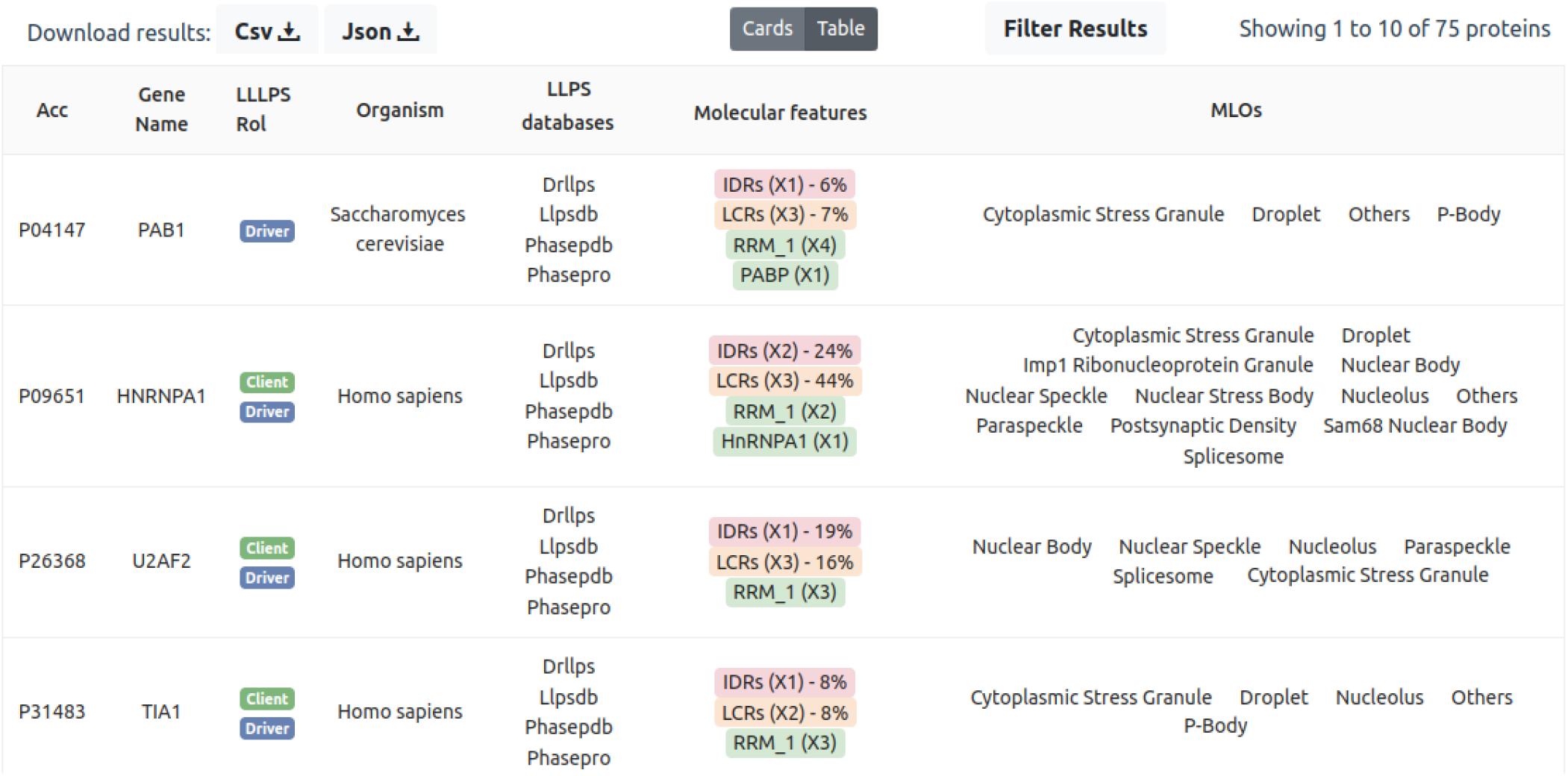
Results page. Example of results page for a query of driver proteins containing RRM_1 domains. Each record shows information related to the LLPS role, LLPS dataset source, Molecular features (Phase separation, Intrinsically disordered regions, Low complexity regions) and MLOs where the protein can be found.

**Figure 4.**
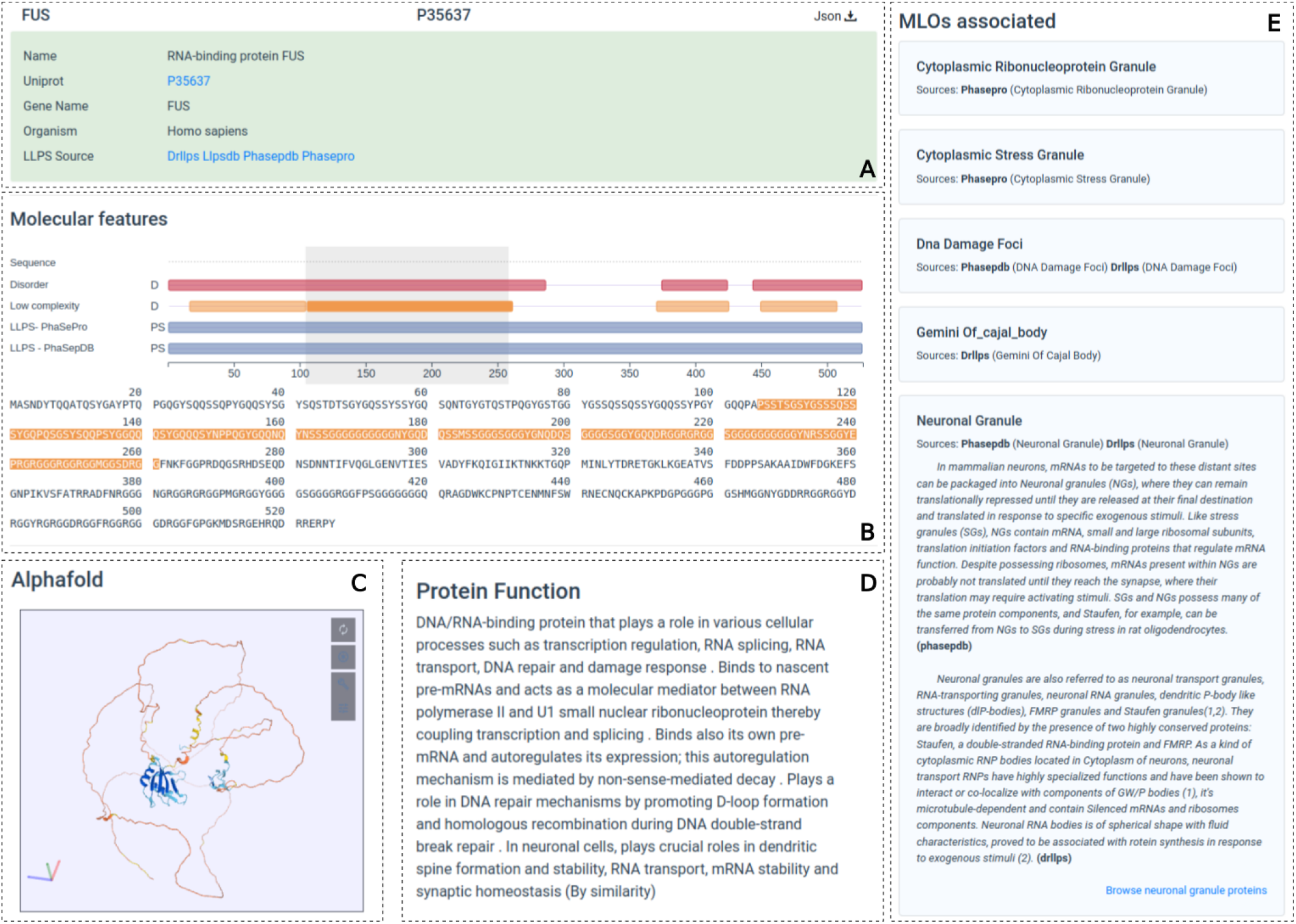
Protein page. A) Protein information and related links to the source databases. B) Molecular features, interactive visualizations of LLPS related molecular features (Phase separation promoting regions, IDRs, LCRs, Prion-like domains, PFAM domains) .C) AlphaFold2 predicted protein structure visualization. D) Protein function (source UniProt). D) Protein biological function. E) Membraneless Organelles associated with the protein. Each MLO has information related to the source database, a brief description of the MLO when available and a link to directly browse proteins associated with the MLO.

In the **protein page**, users will find detailed information of the protein as its sequence, protein and gene names, biological function (when available), LLPS resource database (crosslinked), an interactive feature viewer and sequence viewer with molecular features. It also provides a list of MLOs where the protein is associated. Users will also find the predicted structure by AlphaFold2 with an interactive visualization. For Drivers proteins, a table of Orthologs is provided. All the information is downloadable in JSON format.

In addition, MLOsMetaDB offers an API that allows users to programmatically query and download data, enabling complex searches and facilitating integration into analysis pipelines. The API is thoroughly documented, with detailed explanations of available paths and parameters. Swagger UI is used as a tool for testing queries and results.

## Discussion

Currently, there are few primary databases associated with LLPS and MLO with differences in methodology, objectives, level of curation, number of proteins, and non-uniform, poor, or missing annotation for a limited number of proteins, making further analysis difficult.

In response, MLOsMetaDB offers the scientific community centralized and unified information and predictions on LLPS and MLO-associated proteins, supporting both simple and complex searches. The platform also provides a modern, interactive website for ease of use by non-bioinformatics users. Moreover, MLOsMetaDB includes an API that integrates access, search, and download of LLPS/MLOs protein information into a pipeline, streamlining analyses.

## Abbreviations

API: Application Programming Interface
IDR: Intrinsically disordered regions
LCR: Low complexity regions
LLPS: Liquid-liquid phase separation
MLOs: Membraneless organelles

